# Human organoid modeling of congenital malformations caused by RFX6 mutations reveal an essential role for this transcription factor in establishing and maintaining duodenal identity upstream of PDX1

**DOI:** 10.1101/2023.11.09.566480

**Authors:** J. Guillermo Sanchez, Scott Rankin, Emily Paul, Heather A. McCauley, Daniel O. Kechele, Jacob R. Enriquez, Nana-Hawa Jones, Siri AW Greeley, Lisa Letourneau-Friedberg, Aaron M. Zorn, Mansa Krishnamurthy, James M. Wells

**Affiliations:** Division of Developmental Biology, University of North Carolina at Chapel Hill School of Medicine, Chapel Hill, NC, USA; Center for Stem Cell and Organoid Medicine (CuSTOM), University of North Carolina at Chapel Hill School of Medicine, Chapel Hill, NC, USA; Department of Cell Biology and Physiology, University of North Carolina at Chapel Hill School of Medicine, Chapel Hill, NC, USA; Division of Endocrinology, Cincinnati Children’s Hospital Medical Center, Cincinnati, OH, USA; Division of endocrinology, University of Chicago, Chicago, IL, USA

**Keywords:** Endoderm patterning, Small Intestine, Mitchell-Riley Syndrome

## Abstract

The gastrointestinal (GI) tract consists of highly specialized organs from the proximal esophagus to the distal colon, each with unique functions. Rare congenital malformations of the GI tract, including organ atresia, agenesis or mis-patterning are linked to gene mutations although the molecular basis of these malformations has been poorly studied due to lack of model systems to study human development. We identified a patient with compound heterozygous mutations in the transcription factor RFX6 with pancreatic agenesis as previously described. In addition, the patient had duodenal mal-rotation and atresia suggesting that establishment of the proximal small intestine was impaired in these patients. To identify the molecular basis of the intestinal malformation we generated induced pluripotent stem cell lines from this patient, and derived human intestinal organoid (HIOs) to identify how mutations in RFX6 impact intestinal patterning and function. We identified that the duodenal identity of HIOs and patient tissues had adopted a more distal small intestinal signature, including expression of SATB2, normally expressed in the ileum and colon. CRISPR-mediated correction of RFX6 restored duodenal identity, including expression of PDX1, which is required for duodenal development. Using transcriptomic approaches in HIOs and Xenopus embryos we identified that PDX1 is a downstream transcriptional target of RFX6 and that PDX1 expression in a RFX6 mutant background was sufficient to rescue duodenal identity. However, RFX6 had a PDX1-independent role in regulating expression of components of WNT, HH, and BMP signaling pathways that are critical for establishing early regional identity in the GI tract. In summary, we have identified that RFX6 is one of the most upstream regulators early intestinal patterning in vertebrates and that it acts by regulating key transcriptional and signaling pathways.

## Introduction

Deciphering what factors control specific regional patterning during endoderm organogenesis remains a major area of investigation in developmental biology(; C.A. Thompson, DeLaForest, and Battle 2018; Spence, Lauf, and Shroyer 2011). The mechanisms that drive specific cell migration and identity require specialized gene regulatory networks at precise times to give rise to diverse organs which are regulated by a combination of signaling pathways and transcription factors. Different major signaling pathways such as WNT, BMP and Hedgehog, have been shown to have a critical role in the patterning and formation of endoderm organs (Zorn and Wells 2007, 2009; Roberts et al. 1995; C.A. Thompson, DeLaForest, and Battle 2018). Transcription factors are often responsible for driving organ specific gene regulatory networks and their loss can result in organ agenesis or abnormal patterning. Regulatory factor 6 (RFX6) is a winged-helix transcription factor that has been shown to have an essential role in endoderm organogenesis and is required for endocrine cell formation in pancreatic islets and enteroendocrine cell differentiation in the GI tract (Smith et al. 2010; Piccand et al. 2014). Early in embryonic development RFX6 is broadly expressed in the epithelium of the foregut and proximal intestine and later becomes restricted to the endocrine lineages of the pancreas and the proximal small intestine at later stages of development. Reduction or loss of Rfx6 in mouse intestine resulted in gut malrotation, duodenal atresia, malabsorption, and significant reduction of the enteroendocrine lineage (Soyer et al. 2010; Gehart et al. 2019). In humans, mutations in RFX6 result in Mitchell-Riley syndrome, an autosomal-recessive syndrome of neonatal diabetes, small bowel atresia, and in many cases malabsorption (Kambal et al. 2019; Concepcion et al. 2014; Trott et al. 2020). However, the molecular mechanisms by which RFX6 regulates the patterning and function of the regions of the small intestine remains unknown.

Another transcription factor that has a critical role in the development of the pancreas and proximal small intestine is pancreas/duodenal homeobox 1 (PDX1). It has been shown that RFX6 and PDX1 are co-expressed in the developing duodenum and endocrine cells in pancreas and small intestine (Soyer et al. 2010; Smith et al. 2010; Yang et al. 2017). PDX1 has an essential role in pancreas, stomach and duodenum morphogenesis and persists in the adult tissues, Pdx1-null mouse models have shown a lack of mature pancreatic tissue and malformation of the gastroduodenal junction with significantly reduced endocrine cells in the small intestine with other duodenal function defects (Offield et al. 1996; Boyer et al. 2006; Burlison et al. 2008; Chen et al. 2009; Fujita et al. 2008; Fujitani et al. 2006). In the intestinal tract PDX1 is maximally expressed in the anterior duodenal region with decreased expression in the distal small intestine. Additionally, PDX1 has been reported to be necessary for the production of GIP-secreting cells in the small intestine, although multiple cell lineages are lost in PDX1-null models (Chen et al. 2009; Jepeal et al. 2005). Despite an essential role for PDX1 in regulating patterning and development of the duodenum, it is not known what regulates PDX1 expression.

Over the past decade, several human organoid models have been developed that enable novel strategies to model diseases modeling and organogenesis (Kechele and Wells 2019). Induced Pluripotent Stem Cell (iPSC)-derived organoids are three-dimensional structures that mimic the development of an organ and offer a unique way to study developmental mechanisms and organ functions in human tissue (Carpenter and Rao 2015). Organoids also possess the unique advantage of being generated from iPSCs, meaning they exhibit the same mutation as the patient from whom they are derived, allowing for study of patient-specific phenotypes at multiple different stages of development. iPSC-derived intestinal organoids (HIOs) can be differentiated *in vitro* and engrafted under the mouse kidney capsule where they become vascularized and mature to mimic fetal intestine with crypt-villus architecture and the ability to absorb nutrients (Spence et al. 2011; Watson et al. 2014). We recently used these iPSC-derived human organoids to identify novel patient pathologies in patients with a PDX1 mutation (Krishnamurthy et al. 2022).

We have identified a patient with a compound heterozygous mutation in RFX6 which caused neonatal diabetes and duodenal mal-rotation and atresia. These mutations resulted in functional loss of the RFX6 protein and abnormal gut patterning. We generated iPSCs and HIOs from this patient to further interrogate the specific roles of RFX6 during intestinal development. This approach allowed us to directly compare the patient’s biopsy samples with the HIOs to discover novel pathophysiological effects of this mutation. Here, we demonstrate that RFX6 is required to maintain duodenal identity and function, and that its loss results in ileal-like tissue. We describe the mechanism by which RFX6 turns on PDX1 expression which drives the gene regulatory network that establishes and maintains duodenal identity. Finally, we largely rescue the phenotypes caused by the RFX6 mutations using both CRISPR allele correction and inducible reintroduction of RFX6 or downstream PDX1. This study represents a novel approach to use complex patient pathologies to uncover the mechanisms at play in human development.

## Results

### RFX6 is expressed in the human small intestine and its loss of function leads to gut mispatterning

Several signaling pathways and transcription factors converge to pattern the different segments of the small intestine into duodenum, jejunum, and ileum. These segments carry out different functions during nutrient absorption but importantly, also employ differential gene regulatory networks to generate their identities (Haber et al. 2017). One key transcription factor is the winged helix factor RFX6. Early in development, RFX6 is expressed widely throughout the endoderm but becomes restricted to proximal small intestine and pancreas after E13.5 (Soyer et al. 2010; Smith et al. 2010). A patient presented to CCHMC with monogenic diabetes, duodenal atresia, intestinal malrotation, annular pancreas and the growth of polyps in the duodenum that were surgically removed for analysis. Exome sequencing of the patient and parents revealed that this patient carried a compound heterozygous mutation in RFX6, with a frameshift mutation (p. Arg347Lysfs*18) in the paternal allele and a nonsense mutation (p. Gln875*) in the maternal allele (Fig. 1A). Analysis of resected polyps showed them to be largely gastric in identity expressing the stomach marker CLDN18 with only small regions expressing intestinal marker CDH17 (Fig 1B). This developmental phenotype suggested that early stages of organ development were impacted, for example regional patterning of the gut tube. However, the limited access to patient samples made it challenging to perform deep mechanistic investigation into the cause of these phenotypes. To perform a deep phenotyping and investigate the developmental basis of these congenital malformations we recruited this patient into our study and sent a blood sample to our Pluripotent Stem Cell Facility for generation of induced pluripotent stem cell lines. Analysis of these lines showed them to be karyotypically normal, pathogen free and multipotent (data not shown).

**Fig 1.**
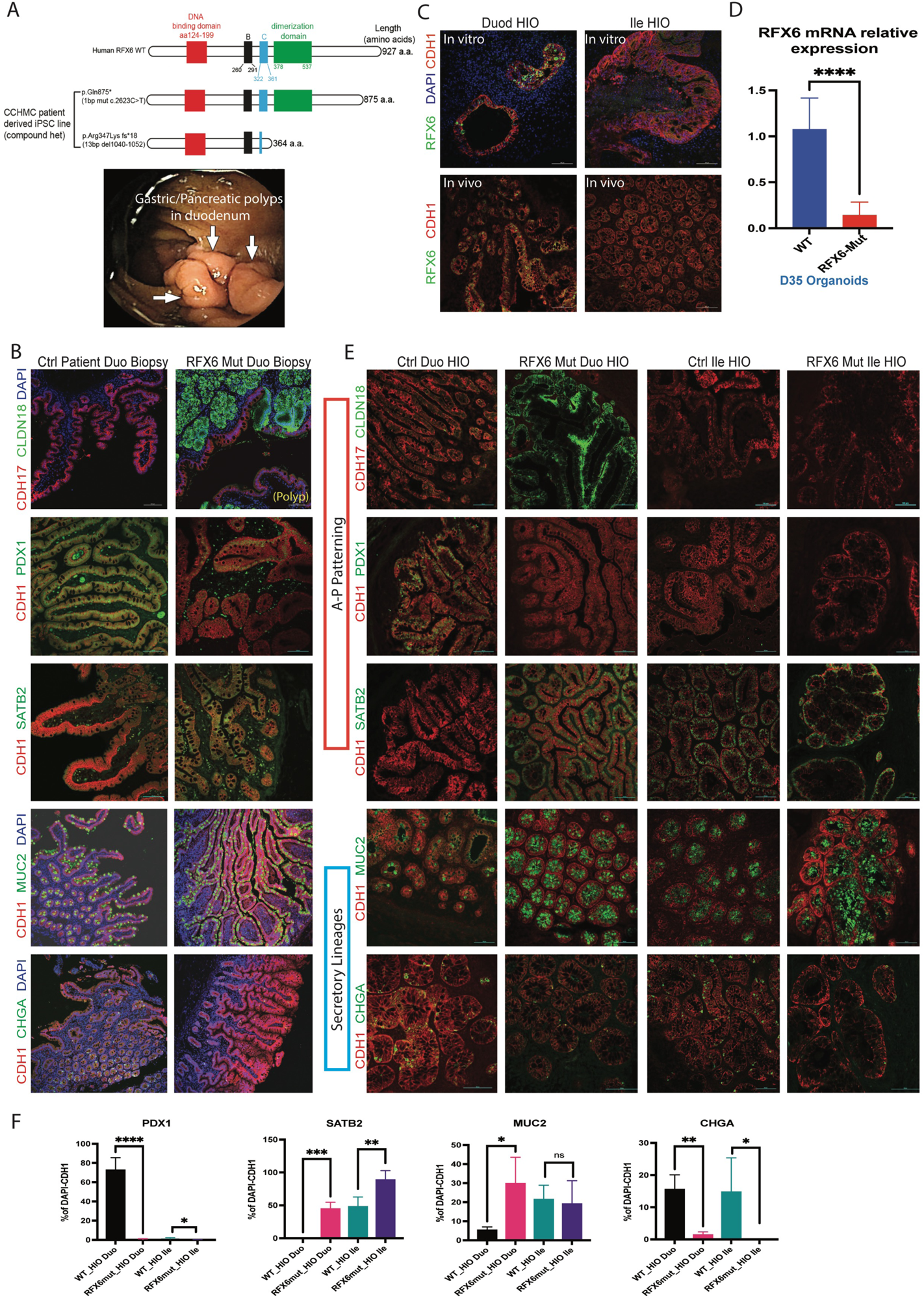
RFX6 mutations cause abnormal gut patterning and loss of endocrine cells in the intestine. A) Patient mutation diagram in paternal and maternal alleles compared to WT allele, endoscopy image of polyps in Duodenum. B) Staining of Rfx6 Mut Patient biopsy compared to healthy duodenal biopsy showing gastric metaplasia (CLDN18), PDX1, SATB2 and MUC2 changes and loss of EECs (CHGA). C) Expression of Rfx6 in Duodenal and Ileal HIOs in vitro and in vivo (RFX6=Green) (scale bar =100um). D) Rfx6 mRNA expression in WT vs Rfx6 Mut HIOs (n=3). E) Staining of Duod and Ileum HIOs from Rfx6 Mut iPSC vs WT Duod and Ileum iPSC HIO shows duodenum phenotypes while ileum is normal excluding loss of EECs. F) Quantification of the staining of the HIOs for the patterning and secretory markers (n=3-8). (markers from panel C) (scale bar=100um). F) Quantification of HIO staining of the Duodenum WT vs RFX6 Mut HIOs (n=3-6). Significance determined by unpaired t-test with *p<0.05, **p<0.01, ***p<0.001.

We first determined if we could phenocopy the patient pathologies with intestinal(duodenal) organoids generated from the patient iPSC lines. We first generated HIOs in vitro, at which stage they represent first trimester human duodenum(Spence et al. 2011). To promote further development and maturation, HIOs were transplanted into immunocompromised mice and allowed to develop for an additional 12 weeks, at which time they are similar to early third trimester human duodenum (Fig. S1A) (Singh et al. 2023). Analysis of human intestinal organoids (HIOs) generated from a control (WT) iPSC line confirmed them to be duodenal in nature, as we previously reported (Spence et al. 2011), expressing PDX1 and CDX2 and little-to-no expression of gastric markers (Fig. S1 B-E). We also found that RFX6 protein was expressed in duodenal HIOs in vitro and those matured in vivo (Fig1C, Fig. S2B-D). In contrast, HIOs made from patient iPSC lines with RFX6 mutations (RFX6 Mut) had low RFX6 expression (Fig. 1D).

Additionally, RFX6 Mut HIOs had large patches of gastric tissue expressing CLDN18, reminiscent of the gastric polyps found in the patient. Further phenotyping of RFX6 Mut HIOs showed a wide range of abnormalities including increased goblet cells and a decrease in enteroendocrine cells with loss of many subtypes (Fig1B, E, S3A). Interestingly, the patient HIOs exhibited a loss of PDX1 expression which is a key driver of the proximal intestine gene regulatory network and an established marker of duodenum (Boyer et al. 2006; Chen et al. 2009). Furthermore, duodenal HIOs had inappropriate expression of the distal intestinal marker SATB2, a transcription factor that has been recently published as a driver of ileal-colonic fate (Fig. 1E) (Gu et al. 2022). As a control for this regional phenotype we generated distal HIOs that are ileal in nature via FGF and Chiron further pattern the developing hindgut (Tsai et al. 2017; Dessimoz et al. 2006). RFX6 is not normally expressed in the distal small intestine, and as predicted distal small intestinal organoids are unaffected by RFX6 mutations (Fig. 1E).

Having identified multiple unappreciated pathologies in RFX6 Mut HIOs, we wanted to investigate if these were similarly found in the patient. Given the pathologic nature of the duodenal polyps, we chose to histologically compare HIOs to a biopsy from a visually normal region of the duodenum from this patient. In all cases, the HIOs phenotypes were also observed in the patient biopsy, including reduced PDX1, increased SATB2, increased goblet cells, and a reduction in most EEC populations (Fig. S3B-D). This confirmed previous murine studies that had reported loss of enteroendocrine cells in Rfx6-null mice (Gehart et al. 2019; Piccand et al. 2019). These data suggest that RFX6 mutations caused mis-patterning of the developing GI tract, with loss of duodenal identity and inappropriate expression of distal markers.

### Transcriptomic analysis reveals abnormal patterning and loss of duodenal function in mature RFX6 patient organoids

To more comprehensively investigate the impact of the RFX6 mutations on intestinal development, we isolated tissue from transplanted HIOs and submitted them for RNA-sequencing. Principle component analysis (PCA) of transcriptomes from WT and RFX6-patient duodenal HIOs showed over 2000 significantly differentially expressed genes between the 2 duodenal samples (Fig. S4A). As expected from our immunofluorescence analysis, genes notably reduced in RFX6 mutant duodenal HIOs included transcripts for several proximally enriched EEC-peptides such as GIP, Ghrelin and Motilin (Fig. S4B), suggesting RFX6 has an important role in the differentiation of intestinal enteroendocrine cells in humans (Piccand et al. 2019).

In addition, we found altered expression of Hox genes, which regulate a plethora of downstream targets to define the anterior-posterior (AP) patterning in developing embryos of species ranging from flies to humans (Fig. S4C)(Hajirnis and Mishra 2021). In the GI tract, HOX genes are expressed in the mesenchyme, the epithelium or both, and follow a proximal to distal gradient going from the duodenum to the colon. In RFX6 Mut duodenal HIOs, distal HOX genes such as HOXC10, HOXA10 and HOXA11 were increased in expression compared to WT duodenal HIOs, suggesting that the duodenum of the RFX6 patient exhibits more ileal identity. However, known proximal HOX genes did not decrease in RFX6 patient duodenal HIOs, suggesting that there is a mix of duodenal and ileal-like gene expression patterns in RFX6-patient HIOs. Gene Ontology (GO) Terms analysis of all the genes downregulated in RFX6 patient duodenal HIOs revealed loss of biological processes associated with duodenal-specific functions (Fig2B). For example. The duodenum is a key segment of the small intestine for lipid uptake and metabolism (DiPatrizio and Piomelli 2015). RFX6-patient duodenal HIOs showed decreased expression of MTTP, APOC3 and RBP2 that are involved in lipid metabolism processes compared to WT duodenal HIOs (Fig. 2A) (Iqbal and Hussain 2009; Hussain 2014). We observed a trend toward increased expression of genes involved in ileal-specific functions such as bile acid absorption and vitamin B12 processing (Fig. 2A)(Ticho et al. 2019; W.G. Thompson and Wrathell 1977).

**Fig 2.**
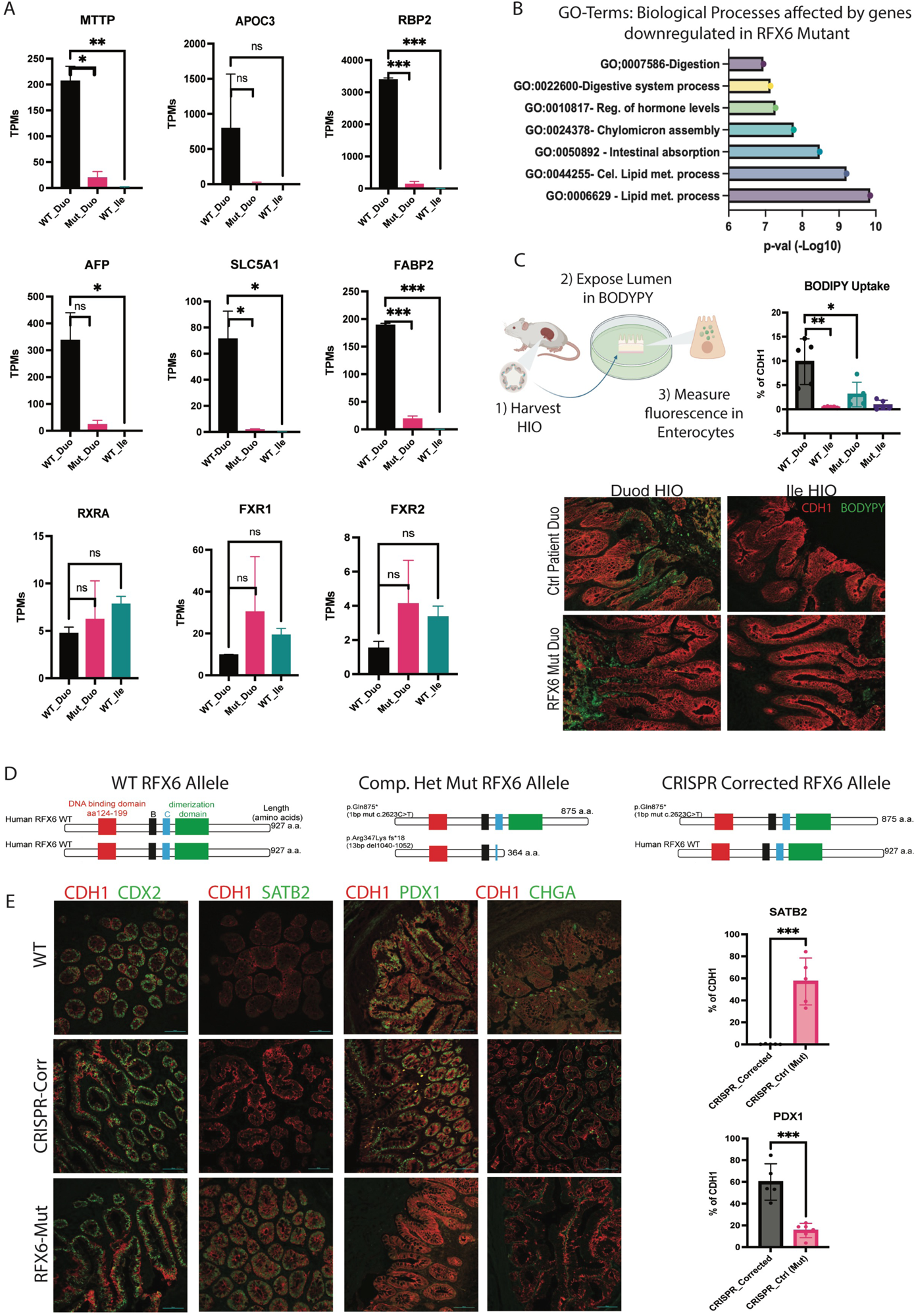
Transcriptomic analysis of RFX6 Mut HIOs shows loss of duodenal function and patterning and can be corrected with one WT allele. A) Genes involved in Duod functions are downregulated in Mut HIOs while genes involved in Ileal functions do not change. Results were obtained from independent transplanted organoids (n=2). B) Gene Ontology results for Biological Processes by genes significantly Downregulated in Rfx6 Mutant HIOs as compared to WT. C) Fluorescently labeled lipid (Bodipy) uptake in WT HIO vs Mut HIOs (n=3). D) Schematic of WT RFX6 allele vs Compound Heterozygous Mutant RFX6 allele vs CRISPR corrected RFX6 Allele. E) Staining of Corrected vs Mutant HIOs compared to WT HIOs showing patterning markers return to normal and Enteroendocrine population is recovered (n=3-7) (scale bar=100um). Significance determined by unpaired t-test with *p<0.05, **p<0.01, ***p<0.001. Significant Differentially expressed genes from Bulk RNA sequencing was defined by adjusted p-value of <.05.

To investigate the impact of RFX6 mutations on lipid absorptive function in RFX6-derived duodenum, we exposed the lumen of organoids to the fluorescent lipid BODIPY and quantified lipid uptake (Carten, Bradford, and Farber 2011). We fist compared BODIPY uptake in WT duodenal HIOs as compared to WT ileal HIOs and as expected observed robust uptake of BODIPY into the epithelium of duodenal but not WT ileal HIOs. In contrast, RFX6 Mut duodenal HIOs did not efficiently absorb lipids (Fig. 2C). Quantification of lipid uptake showed a 2-3-fold reduction in RFX6 Mut duodenal HIOs as compared to control HIOs. Taken together, this data suggests that the RFX6-mutations affects A-P patterning during development of the duodenum resulting in reduction of proximal small intestine functions like lipid absorption and an enrichment of an ileal signature.

### CRISPR correction of one allele was sufficient to restore a more duodenal phenotype in Rfx6 patient organoids

The patient in this study carries compound heterozygous mutations in both parental alleles, with the paternal mutation (p. Arg347Lysfs*18) introducing a premature stop codon that deletes 563 amino acids from RFX6. The maternal allele is a nonsense mutation that deletes 52 amino acids. Because the patient’s parents did not present with clinical manifestation of RFX6 deficiency, we reasoned that conversion of one mutant allele to a WT allele would be sufficient to restore a more normal duodenal phenotype. Previous reports of RFX6 mutations have shown that the larger the deletion, the more severe the symptoms (Kambal et al. 2019). We therefore used CRISPR-Cas9 to convert the paternal mutation into a WT RFX6 allele (Fig 2D) (Haeussler et al. 2016; Ran et al. 2013; Liang et al. 2017). RNA-seq analysis of the transplanted HIOs revealed that the corrected duodenal HIOs were much more similar to the WT duodenal HIOs as compared to RFX6 Mut duodenal HIOs (Fig. S4D-E). Principle component analysis showed that the primary difference caused by RFX6 mutations (PC1;76%) had been completely reversed by CRISPR correction of the paternal allele. However, PC2 showed that there are still differences between control lines and CRISPR-corrected lines. For example, several genes and biological processes specific to the proximal small intestine showed different levels of expression between the WT and corrected HIOs. This could be due to the different genetic backgrounds of the WT control line, or due to reduced RFX6 activity due to the maternal mutation.

A closer analysis of specific factors that regulate duodenal patterning show that expression levels of PDX1, GATA4 and several regionally expressed HOX genes is restored in CRISPR-corrected HIOs (Fig. S4F). We also saw restored expression of genes involved in duodenal function including lipid metabolism (Fig. S4F). CRISPR-corrected Duodenal HIOs did not exhibit evidence of gastric polyps, had significantly decreased expression of ileal-colonic marker SATB2, and goblet cell numbers more comparable to wild-type duodenal organoids (Fig. 2E). Importantly, CRISPR-correction of the paternal RFX6 allele was sufficient to restore the production of enteroendocrine cells (EECs) including GIP, GHRELIN, and MOTILIN producing EECs, populations that are completely lost in the patient derived mutant duodenal HIOs (Fig2E). Together, these data show that the rescue of the paternal allele significantly restored a duodenal phenotype to RFX6 compound heterozygous HIOs suggesting that RFX6 has a critical role in establishing duodenal identity and function. However, the expression of some genes was not restored to the levels of control HIOs generated from iPSC lines of different genetic background. This could be due to haploinsufficiency or gene expression variability between organoids generated from diverse genetic backgrounds.

### Initiation of duodenal patterning and activity of patterning pathways is deranged in RFX6 mutant endoderm

The anterior-posterior regional identity (patterning) of the developing gut tube is established at early stages of development. To investigate the mechanisms leading to the abnormal patterning in RFX6 patient intestine w performed RNAseq analysis during the gut tube patterning stage of differentiation (D7) (Fig. 3A)(Spence et al. 2011; Spence, Lauf, and Shroyer 2011). This stage closely resembles embryonic stages ∼E8.5-9.5 in mice at which point Rfx6 is already expressed (Soyer et al. 2010; Smith et al. 2010). We first confirmed that RFX6 expression in human cultures initiates in this window of time and found that expression starts at D5 and increases through D7 of differentiation (Fig. S2B), PCA analysis of RNAseq data displayed a 94% variance between WT and RFX6 Mut and over 4,000 differentially expressed genes (Fig. 3B, S5A). Most of the differentially expressed genes were downregulated in the RFX6-Mut patient line, suggesting a failure to activate an RFX6-mediated transcriptional program (Fig. S5A). Lineage tracing experiments in mice show that RFX6 expression is restricted to the epithelium at this stage of development (Smith et al. 2010). Consistent with this we observed that critical endodermal patterning regulators like CDX2, GATA4, GATA6, HNF4A were significantly reduced in the D7 gut tube endoderm cultures from RFX6 mutant iPSC lines (Fig. 3C). Consistent with our observation that the patient duodenum had areas of mixed organ identity, we used a list of organ markers from a published atlas and found that RFX6 Mutant cultures lost small intestinal organ markers such as ling and stomach (Fig. 3D) (Yu et al. 2021). In addition to epithelial patterning defects, we observed a surprising number of expression differences in key signaling factors that regulate endodermal patterning through a series of epithelial-mesenchymal interactions (Zorn and Wells 2007; Walton and Gumucio 2021; McCarthy et al. 2020) (Fig. 3C). In particular, WNT3, WNT5B, BMP4, IHH and many downstream targets of these pathways were reduced in RFX6 mutant cultures. Since WNT signaling is involved in the expression of CDX2, the master regulator of intestinal identity(Sherwood et al. 2011), it is likely that reduced WNT signaling in D7 cultures causes reduced CDX2 expression. Analysis of known CDX2 targets that are involved in early intestinal development (Kumar et al. 2019) are also reduced in the RFX6-Mut patient line suggesting that the abnormal patterning is due to low CDX2 expression (Fig. S5B). While RFX6 is expressed in the epithelium, RFX6 mutations cause patterning changes in both the epithelium and mesenchyme of D7 cultures. To begin to separate the direct effects of the RFX6 mutations in the epithelium versus the indirect effects on the mesenchyme, we looked at published dataset from Han et al. which mapped endoderm vs mesoderm specific markers in early gut development (Han et al. 2020). We identified endoderm and mesoderm specific markers and performed Gene Ontology analysis. Interestingly, the biological processes that were most enriched for this common genes lost in our RFX6 Mutant cultures included intestinal patterning, epithelium development, tube development and morphogenesis (FigS5C-D). Together, these data suggest that there is an initial delay of intestinal patterning as observed by low CDX2 expression in RFX6 mutant cultures. However, at later stages of differentiation, CDX2 is expressed in both control and RFX6 mutant HIOs and at those stages duodenal markers are lost and more distal markers are expressed.

**Fig 3.**
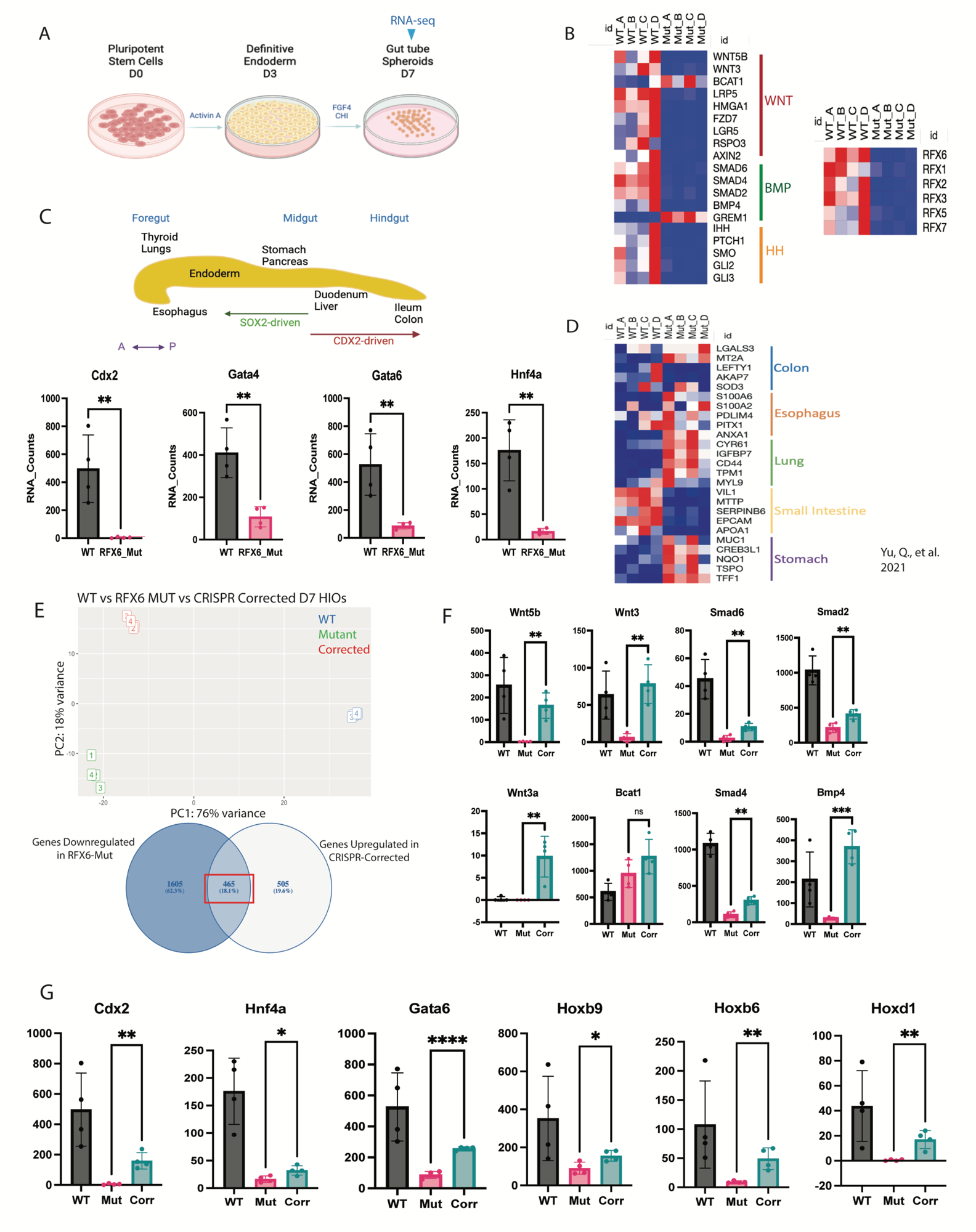
RFX6 Mut D7 HIOs show mis-regulation of signaling pathways and early hindgut patterning, corrected with one WT allele. A) Schematic of Organoid generation and harvesting timepoint for bulk RNA sequencing. B) Changes in components of relevant signaling pathways that are affected in Rfx6 Mut D7 HIOs (Wnt, BMP and HH pathways) and other members of the RFX family based on RNA Counts (n=3). C) Transcription factors involved in mid/hindgut patterning are also affected in Rfx6 Mut D7 HIOs based on RNA Counts (n=3). D) Organ markers from atlas showing intestinal markers are down in Rfx6-Mut HIOs based on RNA Counts (n=3) (Yu et al. 2022) E) PCA of WT vs RFx6 Mut and CRISPR Corrected D7 HIOs with percentage of genes upregulated by CRISPR Correction, 18.1%. F) Upregulated components of several signaling pathways increase in CRISPR Corrected D7 HIOs based on RNA Counts (n=3). G) Patterning markers return to levels closer to control in CRISPR Corrected D7 HIOs based on RNA Counts (n=3). Significance determined by unpaired t-test with *p<0.05, **p<0.01, ***p<0.001. Significant Differentially expressed genes from Bulk RNA sequencing was defined by adjusted p-value of <.05.

Finally, we determined if the CRISPR-correction of the paternal allele rescued the patterning of the HIOs at D7 using principle component analyses of RNAseq data. CRISPR correction restored the PC1 component (76% variance) to nearly wildtype, suggesting that correction of one allele allows transcription of many downstream targets. There was, however, an 18% variance in PC2 from the WT and the Corrected, which could be due to the fact that the WT iPSC line comes from a different genetic background (Fig. S5E). Transcripts that were restored in the CRISPR corrected cultures included components of the WNT and BMP pathways known to regulate gut tube patterning (Fig. 3F). Lastly, we found that the correction of one RFX6 allele was sufficient to rescue the expression of several key transcription factors in hindgut development and patterning such as CDX2 and HNF4A (Fig. 3G, S5F). Taken together, this data suggests that while RFX6 expression is specific to the epithelium, it functions to regulate patterning and signaling in both the epithelium and mesenchyme. Additionally, the CRISPR correction of the paternal allele provides evidence show that the maternal mutation has similar levels of activity than a WT allele.

### An evolutionary conserved role of RFX6 in GI tract patterning and functional characterization of RFX6 mutations

Our CRISPR-Correction of one allele in the RFX6 Mut line caused the change in expression of many TFs, specifically, we saw an increase in PDX1 RNA expression which had been lost in the mutant. Additionally, expression of hindgut marker SATB2 decreased with the CRISPR correction (Fig. 4A). But we wanted to understand whether these were direct interactions regulated by RFX6. To investigate if RFX6-dependent GI tract patterning is evolutionarily conserved and to further test the functional impact of the 2 identified mutations in human RFX6, we performed a series of loss-of-function and gain-of-function experiments in the amphibian Xenopus, a vertebrate model in which endoderm development and GI tract organogenesis is conserved (Zorn and Wells 2009; Rankin et al. 2021). Xenopus rfx6 shares a similar expression pattern as observed in mammals (Pearl, Jarikji, and Horb 2011; Smith et al. 2010) which we verified by in-situ hybridization in foregut progenitors at stage NF20 (∼1 day post fertilization; “d.p.f.”) and in pancreas, stomach and duodenum progenitors at NF33 (∼2.5 d.p.f.) (Fig. S6A). Knockdown of rfx6 in xenopus using a previously validated translation-blocking morpholino (MO; (Pearl, Jarikji, and Horb 2011)) reveal a loss of pdx1 at NF33 and microdissection of the gut tube at NF43 (∼4d.p.f.) revealed a hypoplastic, mal-rotated GI tract with strong reduction of the pdx1 stomach/duodenum domain (Fig. S6B), a phenotype similar to the mouse Rfx6-/- knockout (Smith et al. 2010). Interestingly, we observed an expansion of the distal hindgut satb2+ domain into the midgut and foregut territories at NF33, suggesting an evolutionary conserved function for rfx6 in regulating vertebrate GI tract patterning. Injection of mRNA encoding human RFX6 Was able to rescue these molecular changes at NF33 as well as formation of a distinct stomach/duodenum domain at NF43 (Fig. 4B, Fig. S6B).

**Fig 4.**
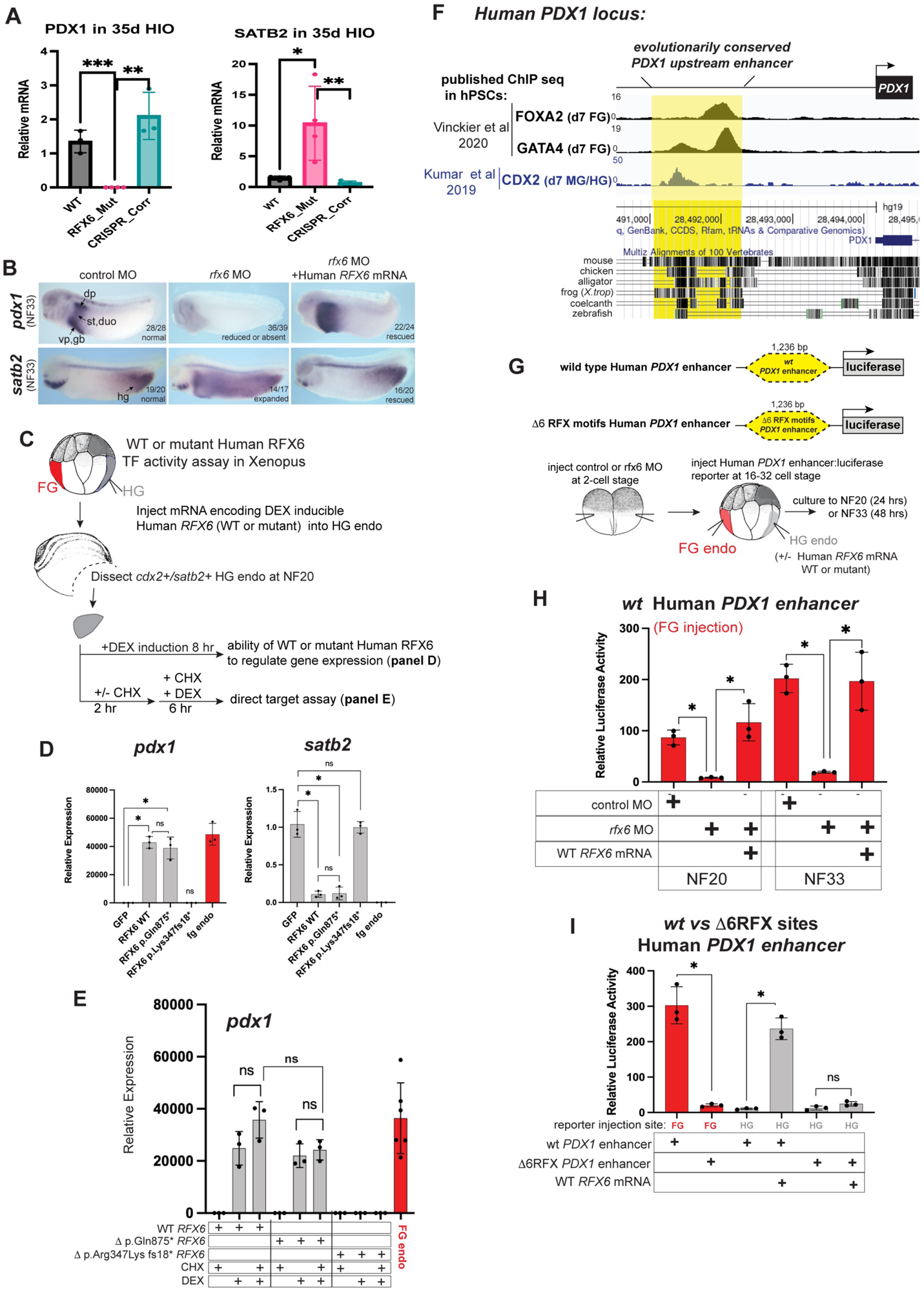
An evolutionarily conserved role of RFX6 in GI tract patterning and functional characterization of RFX6 mutations. A) RT-PCR expression of PDX1 and SATB2 when correcting one allele of RFX6 in 35d HIOs (n=3). B) In situ hybridization of*pdx1* (expression domain abbreviations: st, stomach; duo, duodenum; vp, ventral pancreas; dp, dorsal pancreas; gb, gall bladder) and *satb2* (abbreviation: hg, hindgut) in stage NF33 (2.5 d.p.f) Xenopus embryos. Numbers in the lower left corner of panel indicate number of embryos assayed with the shown gene expression pattern. C-E) Experimental schematic and human RFX6 transcription factor (TF) activity assay in Xenopus. Dexamethasone (DEX) inducible expression constructs were generated for human RFX6 WT, p.Gln875*stop, and p.Arg347Lys.fs18*stop as described in the text. mRNA encoding each was in vitro transcribed and then injected and pools of n=4 explants were pooled in triplicate for RT-qPCR analysis of gene expression, normalized to the housekeeping gene *odc* and relative to uninjected, untreated HG endo explants (D-E). D) RT-qPCR analysis of HG endo explants after 8hours of TF (DEX) induction shows that human WT RFX6 and human p.Gln875*stop. E) *pdx1* is a direct target of RFX6. Red bar= expression of endogenous *pdx1* in foregut endoderm explants (relative to uninjected, untreated HG endo) as a comparison. **F**) Visualization of an evolutionarily conserved upstream *PDX1* enhancer. UCSC genome browser shot of the human PDX1 locus is shown below with tracks from the indicated species, with black bars indicating regions of evolutionary conservation. (**G**) Experimental schematic of the Xenopus luciferase reporter assay used to test RFX6 dependent regulation of the human *PDX1* enhancer. (**H-I)** RFX6 regulates human *PDX1* enhancer activity and mutation of 6 CisBP predicted (Weirach et al 2014) RFX motifs abolishes RFX6 responsiveness. Each black dot in the luciferase activity graphs represents pool of n=5 embryos, mean relative luciferase activity +/- standard deviation is graphed. Asterisks*= p<0.05, parametric two-tailed paired t-test.

We next utilized the Xenopus model to further interrogate the functional impact of the two observed mutations in human RFX6. We constructed Dexamethasone (DEX) inducible version of human WT RFX6 and the 2 mutants, in which the ligand-binding domain of the human glucocorticoid receptor (GR) is fused to the N-terminus of RFX6;this GR-Dex induction system has been widely used for decades in Xenopus to temporally control transcription factor activity (Sive and Bradley 1996; Zorn, Butler, and Gurdon 1999; Rankin et al. 2021). mRNA encoding the GR-TF fusion is synthesized in vitro and injected into embryos; the mRNA is translated in vivo by the embryo and the GR domain functionally sequesters the TF in the cytoplasm until addition of DEX to the culture buffer, at which point the TF can translocate to the nucleus and regulate target gene expression.

We compared the ability of human WT RFX6 and the 2 mutants to ectopically induce*pdx1* and suppress *satb2* in targeted injections into the Xenopus hindgut endoderm (Fig.4C), utilizing a hindgut explant dissection assay which also allowed for assessment of direct induction via pre-treatment of explants with cycloheximide (CHX) before temporal DEX TF induction; CHX blocks indirect targets from being translated and therefore genes induced in the presence of CHX are considered immediate direct targets (Fig.4C-E). WT human RFX6 and the p.Gln875*stop mutant were both able to robustly induce *pdx1* and suppress *satb2* in Xenopus hindgut endoderm, whereas the p.Arg347Lys.fs18*stop mutant had no activity (Fig.2D), strongly suggesting this mutation creates a null, non-functional allele. Similarly in the CHX direct target assay, both the WT and p.Gln875*stop mutant were able to induce *pdx1* in the presence of CHX, suggesting that *pdx1* is a direct target of RFX6 in vivo, whereas again the p.Arg347Lys.fs18*stop mutant had not activity (Fig.4E). As a positive control for CHX effectiveness, we assayed the expression of the pancreas TF ptf1a, whose induction by RFX6 was largely blocked by CHX treatment (Fig.S6C). Together these results suggest the p.Gln875*stop mutation, resulting in a loss of the last 52 amino acids of the C-terminus of RFX6, does not affect RFX6 transcriptional activity, whereas the more severe mutation resulting in p.Arg347Lys.fs18*stop creates a null, non-functional allele in vivo.

To further investigate the direct induction of *pdx1* by RFX6 we bioinformatically examined a known, evolutionarily conserved enhancer upstream of human *PDX1,* of which the orthologous mouse region is sufficient drive reporter expression in pancreas, stomach, and duodenum (Gannon, Gamer, and Wright 2001; Fujitani et al. 2006) indicating functional enhancer activity. Evolutionary comparison of this human *PDX1* upstream enhancer region (hg19_dna range=chr13:28491064-28492299) revealed significant conservation amongst mouse, chicken, alligator, frog, coelacanth, and zebrafish genomes; using the TF motif searching tool CisBP (Weirauch et al. 2014), 6 potential RFX binding motifs were predicted (Table S1).). Analysis of published ChIP-seq studies in human PSCs during directed endoderm differentiation for key endoderm identity TFs FOXA2 and GATA4 (Vinckier et al. 2020), as well as for the midgut/hindgut TF CDX2 (Kumar et al. 2019), revealed binding of these important TFs to this conserved human *PDX1* upstream enhancer (Fig.4F), suggesting it also regulates developmental expression of *PDX1* in human endoderm. We thus utilized the Xenopus model to interrogate potential RFX6-dependent regulation of this predicted human *PDX1* enhancer. Luciferase reporters were constructed containing the WT human *PDX1* upstream enhancer region or a version in which all 6 CisBP-predicted RFX motifs were mutated (Fig.4G). Injection of the WT Human PDX1:luc reporter into foregut endoderm (targeting the progenitor domain of the pancreas, stomach, duodenum) revealed robust reporter activity whereas injection of the enhancer:luc reporter into the satb2+ hindgut domain resulted in negligible activity (Fig.4H, I) at both NF20 and NF33. *Endogenous* Xenopus Rfx6 was necessary for reporter activation in the foregut, as Rfx6 knockdown via morpholino prevented foregut targeted *PDX1* enhancer reporter activity. In support of this, ectopic injection of WT human RFX6 mRNA into the satb2+ hindgut domain resulted in robust PDX1 enhancer activity (Fig.4I), and mutation of the 6 CisBP-predicted RFX motifs in the human PDX1 enhancer abolished reporter activity in the foregut domain as well as negated the ability of WT human RFX6 to activate the reporter in the hindgut (Fig.4I). We further verified that the human p.Gln875*stop RFX6 mutant retained ability to activate the human *PDX1* enhancer, whereas the p.Arg347Lys.fs18*stop mutant had no ability to activated the enhancer (Supplemental Fig.S6D). Together these results suggest RFX6 controls vertebrate GI tract pattern in part via direct activation of *PDX1* via a conserved upstream enhancer.

### Identifying PDX1-dependent and independent roles for RFX6 in patterning in D7 gut tube cultures

RFX6 and PDX1 have overlapping expression patterns in the nascent duodenal endoderm and both genes are required for normal duodenal development. Our data shows that PDX1 expression is lost in RFX6 mutants and suggests that PDX1 is a direct RFX6 target. This suggests that some or all on the phenotypes associated with loss of RFX6 could be due to loss of PDX1 expression. To identify PDX1-dependent versus independent roles of RFX6, we used RFX6 mutant iPSCs to generate tetracycline-inducible RFX6 or PDX constructs whereby we could induce expression of either gene at specific timepoints (Fig. S7A-B) and identify downstream transcriptional responses. We differentiated RFX6 mutant iPSCs and induced expression of either RFX6 or PDX1 separately after D4 of differentiation and maintained until harvest at D7, the gut tube stage (Fig. 5A). The PCA revealed that gut tube endoderm expressing RFX6 induction was more similar to the WT than cultures where we induced PDX1 expression (Fig. 5B). A comparison of differentially expressed genes in response to RFX6 or PDX1 expression shows that RFX6 expression rescued 58% of the transcripts that were down in the RFX6 mutant cultures. In contrast tet-induced PDX1 expression rescued only 25%of the transcripts lost in the RFX6 mutant cultures (Fig 5C, S7C-D). From these comparisons we identified which DE genes in the RFX6 mutant gut tube cultures were PDX1-dependent (rescued by PDX1 expression) as compared to those that were not rescued by PDX1 expression. A gene ontology analysis of genes that appear to be in the RFX6-PDX1 regulatory axis involved duodenal development (Fig. 5D) Expression of either RFX6 of PDX1 in the RFX6-mutant line was able to rescue several key gut patterning markers such as CDX2, SATB2, and GATA4, although PDX1 induction was not able to rescue RFX6 expression, supporting that PDX1 acts downstream of RFX6 (Fig. 5E). While the majority of PDX1 regulated genes are also regulated by RFX6, 67% of the 2070 DE genes in the RFX6 mutant samples were not regulated by PDX1, suggesting that these are regulated by RFX6 independently from PDX1 (Fig. 5D, S7E). In particular, RFX6 appears to target expression of numerous genes involved in signal transduction pathways that play essential roles in gut tube patterning and development such as WNT, BMP and Hedgehog. RFX6 significantly upregulated transcription of multiple ligands, antagonists and receptors suggesting that RFX6 impacts intestinal development is through regulation of these important signaling pathways (Fig. 5F). Lastly, we observed that RFX6 induction is also able to upregulate other members of the RFX family, mainly RFX4-7, whereas PDX1 fails to induce most RFX-genes (Fig. 5G). Taken together, these data show that RFX6 is responsible for the correct patterning of the small intestine via the activation of PDX1 in the gut tube epithelium as well as through regulating signaling pathway components that are known to regulate gut tube patterning.

**Fig 5.**
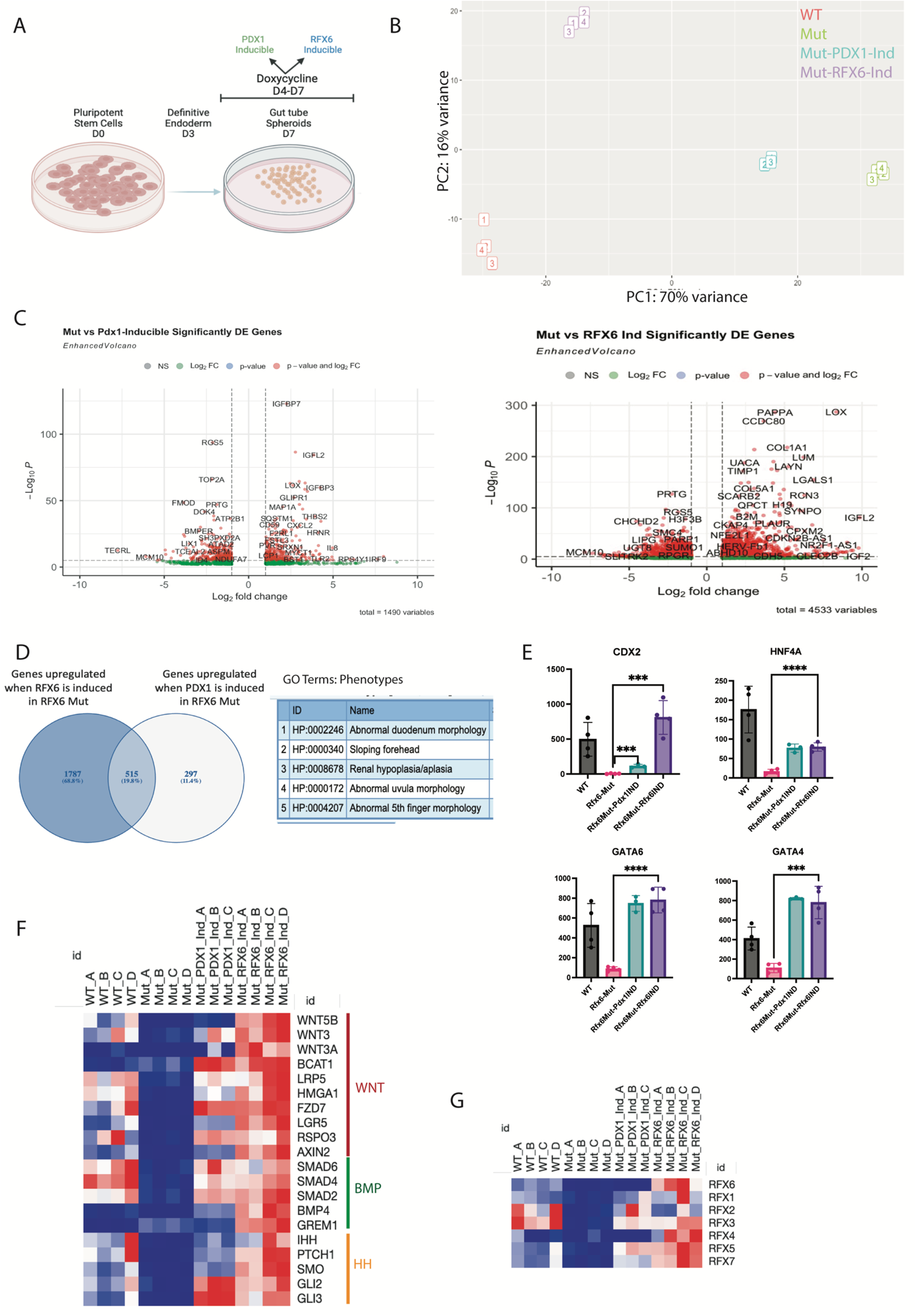
RFX6 and PDX1 inducible constructs rescue early hindgut patterning in RFX6 Mut D7 HIOs. A) Schematic of induction of PDX1 and RFX6 during D4-D7 of HIO differentiation. B) PCA of WT vs RFx6 Mut and PDX1-IND and RFX6-IND D7 HIOs (n=4). C) Genes that are upregulated in RFX6 Mut when inducing PDX1 and RFX6 when compared to RFX6 mutant with no induction. D) Overlap of Genes upregulated by inducing PDX1 and RFX6 in RFX6-Mut D7 HIOs E) Induction of either RFX6 or PDX1 in RFX6 Mut D7 HIOs upregulate expression of TFs that have a role in hindgut patterning F) Induction of RFX6 upregulates signaling pathways in midgut patterning in RFX6 Mut D7 HIOs while induction of PDX1 is less effective. G) RFX6 induction in RFX6 Mut D7 HIOs upregulates expression of other RFX genes. Significance determined by unpaired t-test with *p<0.05, **p<0.01, ***p<0.001. Significant Differentially expressed genes from Bulk RNA sequencing was defined by adjusted p-value of <.05.

### Long-term expression of PDX1 is required to stably maintain duodenal identity in RFX6 mutant HIOs

We identified that PDX1 expression in the RFX6 mutant gut tube endoderm was sufficient to rescue early expression of duodenal patterning markers. However, it was not clear if early rescue of duodenal patterning was stable at later stages of development. To test this, we used the PDX1 doxycycline-inducible line to express PDX1 during the early stages of duodenal development that occurs in vitro and then we extended development for an additional 10 weeks by transplanting HIOs into immunocompromised mice (Fig. 6A). We found that expression of PDX1 during HIO growth *in vitro* was not sufficient to maintain duodenal identity during the added 10 weeks of development in vivo (Fig. 6B-C). Moreover, we observed restoration of transcripts of proximal small intestine specific functions such as lipid metabolism, transport, and absorption (Fig. 6D) including APOB, APOA4 and APOC3 (Fig. 6E), suggesting this phenotype associated with loss of function of RFX6 could be restored by PDX1. However, if we maintained PDX1 expression *in vivo* by feeding transplanted mice with Dox-Chow, we observed restoration of markers of normal duodenal patterning and more normal numbers of goblet cells (Fig. 6F). While we also saw an increase in the number of EECs in PDX1 expressing HIOs, they were still lower in number as compared to controls (Fig. 6F-G). One possible reason is that EECs have higher levels of PDX1 than other epithelial cell types and DOX chow might not induce high enough levels of PDX1 expression to promote robust EEC differentiation. We therefore augmented DOX levels by injecting transplanted mice with an additional 10ug/kg dose of Dox 5 days before harvesting the HIOs. This resulted in rescuing the number of EECs in RFX6 mutant HIOs to nearly WT levels (Fig. 6H). Our fata suggests that PDX1 does not only initiate the patterning of the duodenum and midgut, but it is required to be maintained to keep duodenal function and identity (Fig. 6I). Altogether, these data show that expression of PDX1 can partially rescue the phenotypes seen in RFX6 mutant duodenal HIOs and that PDX1 must be actively maintained to preserve duodenal identity.

**Fig 6.**
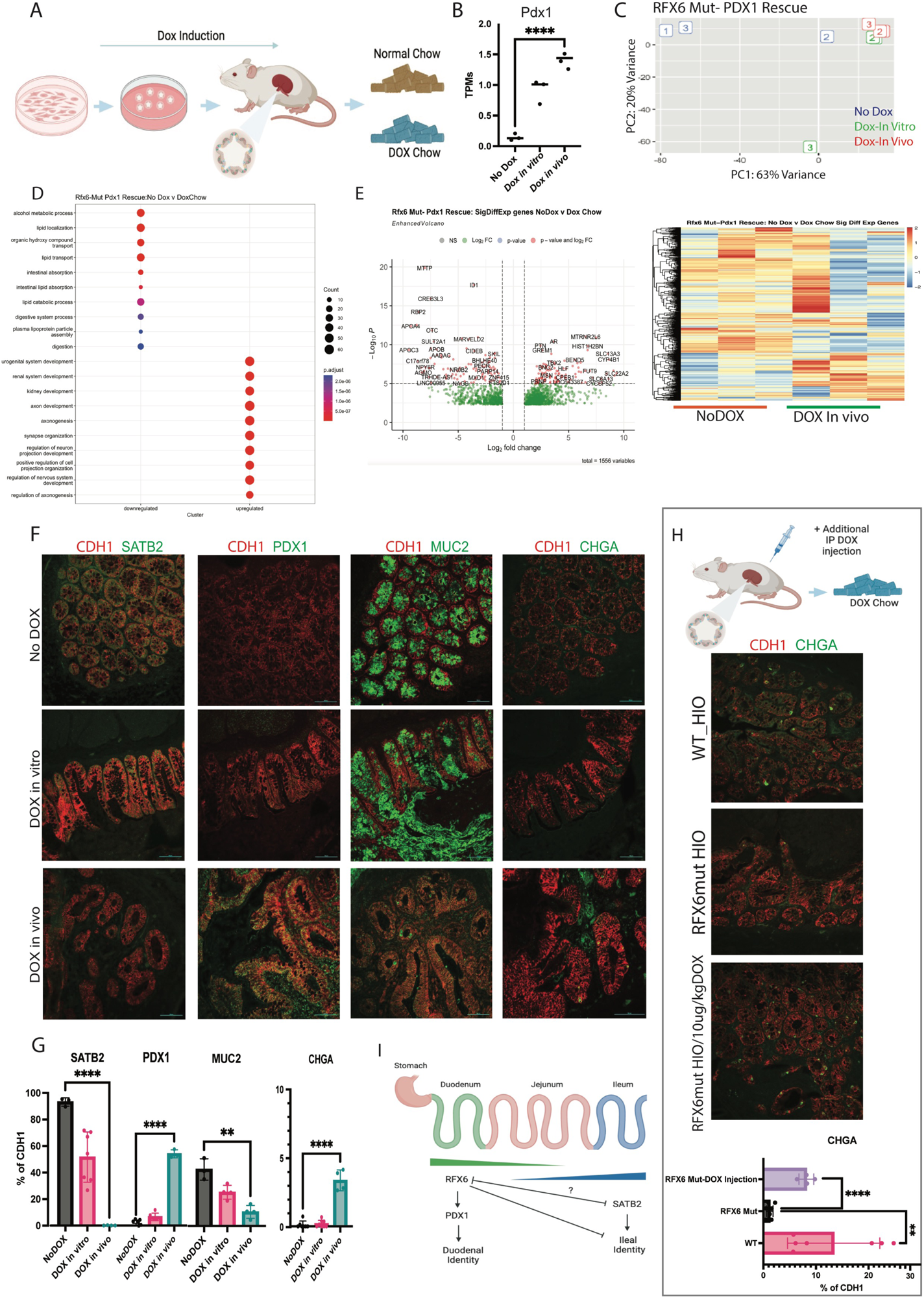
Prolonged induced expression of Pdx1 can partially rescue Rfx6 mutant phenotypes. A) Schematic or Pdx1 induction via Doxycycline in Rfx6 Mut HIOs during the differentiation or*in vivo* via chow. B) PDX1 expression in induced RFX6 mut HIOs after transplant and difference with and without induction *in vitro or in vivo*. C) PCA and Heatmap of RFX6 Mut HIOs with DOX PDX1-inducible in vitro and in vivo (n=3). D) Gene ontology of biological processes shows rescue of Duodenal specific functions in RFX6 Mut HIOs with Dox-induced PDX1 both *in vitro and in vivo*. E) Differentially expressed genes between RFX6 Mut HIO with no PDX1 inducible DOX and DOX in vivo. F) Staining comparison of patterning (SATB2, PDX1) and secretory (MUC2, CHGA) markers in RFX6 Mut HIOs with and without Dox PDX1-Induced in vitro and in vivo (scale bar=100um). G) Quantification of patterning and secretory markers in RFX6 Mut HIOs with and with no DOX treatment (N=3-8). H) Additional injection of Dox to induce PDX1 has a bigger effect in EEC differentiation in RFX6 mut HIOs. I) Proposed mechanism of Rfx6 role in duodenal patterning, acting upstream of PDX1 and repressing distal intestinal features. Significance determined by unpaired t-test with *p<0.05, **p<0.01, ***p<0.001. Significant Differentially expressed genes from Bulk RNA sequencing was defined by adjusted p-value of <.05.

## Discussion

In this study, we used human intestinal organoids (HIOs) to uncover the molecular basis of the intestinal phenotypes observed in a patient Mitchell-Riley Syndrome caused by a compound heterozygous mutation in RFX6. Mitchell-Riley syndrome is a recessive syndrome characterized by neonatal diabetes, pancreatic hypoplasia, intestinal atresia, chronic diarrhea and malrotation(Concepcion et al. 2014). We identified that RFX6 regulates early patterning of the proximal small intestine and the maintenance of duodenal identity and used transcriptomics to identify possible downstream targets. As previously published, pluripotent stem cell-derived HIOs are a tractable system to study the genetics of human GI development (Spence et al. 2011; Krishnamurthy et al. 2022). Genetic correction via CRISPR-Cas9 of one allele of RFX6 was sufficient to correctly pattern the small intestine and maintain duodenal identity. A tetracycline-inducible approach also demonstrated that PDX1 is one of the main functional downstream targets of RFX6 to restore duodenal identity in RFX6-deficient HIOs. We also identified RFX6-dependent, PDX1-independent gene expression changes in numerous regulators of WNT, HH, and BMP signaling pathways that regulate the establishment of regional patterning in the developing gut tube, including the early expression of CDX2. At later stages, RFX6 mutations lead to abnormal A-P patterning measured by loss of duodenal identity and function and acquisition of an ileal program.

RFX6 mutations caused several phenotypes in the intestinal organoids, however, we wanted to see if these could get corrected. CRISPR correction of one RFX6 allele rescued the phenotypes at the gut tube development stage and at later stages of development one WT allele is sufficient to restore intestinal development and maintain duodenal identity. In addition, RFX6 mutants were missing several enteroendocrine cell types that were rescued by correcting the paternal allele. In addition to PDX1 acting downstream of RFX6 during gut tube patterning, PDX1 is co-expressed with RFX6 in enteroendocrine cells (EECs) of the duodenum (Chen et al. 2009; Boyer et al. 2006; Fujita et al. 2008; Fujitani et al. 2006; Yang et al. 2017). Since the doxycycline-inducible construct allowed us to introduce PDX1 at different timepoints, we were able to show that PDX1 induction can correct the intestinal phenotypes caused by the RFX6 mutation as early as the hindgut development stages. At later stages of intestinal development, when RFX6 and PDX1 play essential roles in the differentiation of EEC lineage, we observed that PDX1 expression can rescue the EEC phenotype observed in the RFX6 mutant HIOs. However, this required us to maintain high levels of PDX1 expression throughout all stages of*in vitro* and *in vivo* development. This suggested that EEC development required sustained, high levels of PDX1.

Additionally, we show a significant overlap in differentially expressed genes when inducing RFX6 and PDX1 expression respectively which is evidence for the redundancy in the role of these transcription factors in intestinal maintenance. Finally, we show that PDX1 not only has to be expressed during the gut developing stage but is required to be continually on to maintain duodenal identity. We are able to identify a plethora of phenotypes caused directly and indirectly by loss of RFX6 which suggests that it has a role in regulating the cross-talk between mesenchyme and epithelium during gut development. We are able to identify the epithelial-specific mechanism of duodenum maintenance via PDX1 and identify candidates that could be the intermediate steps in the regulation of signaling pathways between mesenchyme and epithelium. Additionally, along with evidence from both parents normal RFX6 function and evidence from our CRISPR correction and Xenopus data, we are able to determine that the maternal allele retains transcriptional activity while the paternal allele is equivalent to complete loss-of-function. This provides an insight on RFX6 function relating to each domains of the protein. Further studies are needed to deep dive into a structural-functional correlation analysis, however, our data suggests that the paternal allele, having the DNA binding domain intact, still loses nearly all RFX6 transcriptional activity.

As a summary, we have shown that HIOs from patient-derived iPSCs faithfully replicate the phenotypes seen in RFX6 patient biopsies and used this model system to study the role of this transcription factor in human duodenal development. We showed one WT allele of RFX6 is sufficient for normal intestinal development and can also rescue these phenotypes via downstream target PDX1 induction. Some limitations of the study were that there is variability between organoids and specially between parental cell-lines, additionally the differentiation protocol requiring kidney capsule transplantation is technically challenging and lengthy. However, this study helps uncover the role of RFX6 in the development of the intestine and the maintenance of intestinal function with the characterization of a downstream target in PDX1. This study uses technologies that are broadly translatable to other organs which opens the path for engineered tissue to be an alternative for transplantation after genetic correction.

## MATERIALS AND METHODS

### PLURIPOTENT STEM CELL CULTURE AND DIRECTED DIFFERENTIATION OF HIOS

The iPSC lines were generated from the patient as previously described (Zhu et al. 2016). iPSCs were maintained in feeder-free culture. Cells were plated on hESC-qualified Matrigel (BD Biosciences, SJ CA) and maintained at 37C with 5%CO2 with daily removal of differentiated cells and replacement of mTeSR1 media (STEMCELL Technologies, Vancouver Canada). Cells were passaged routinely every 4 days using Dispase (STEMCELL Tec.). HIOs were generated according to protocols established in our lab (Spence et al. 2011; Kechele and Wells 2019; Tsai et al. 2017) and experiments with human iPSCs were approved by the CCHMC ESCRO committee (Protocol #EIPDB2713).

### IN VIVO TRANSPLANT OF HIOS

28-35 days after spheroid generation, HIOs were removed from Matrigel and transplanted under the kidney capsule of immune deficient NOD. Cg-Prkdc^scid^Il2rg^tm1Wjl^/Szj (NSG) mice (Watson et al. 2014). NSG mice were maintained on Bactrim chow for a minimum of 2 weeks prior to transplantation and thereafter for the duration of the experiment (8-14 weeks).

### CRISPR/CAS9 CORRECTION

CRISPR/Cas9 was used to correct the mutation in the paternal allele in the Rfx6 patient iPSC line. The guide RNA used was ACATAAAAATTGGGAACAGT. A phosphorothioated single-stranded DNA donor oligo (T*A*A*TGCATTTTTTAACAATGAAGCATTTAACACATAGCCTTCTTTGTAGCTTATTAGCAGAC<u>ATA</u>AGA AATTTTG<u>CTAAg</u>AAcTGGGAgCAaTGGGTTGTTTCATCCTTGGAAAACTTGCCAG*A*A*G) was designed to include the nucleotide insertion according to published methods(Liang et al. 2017.) The iPSCs were reverse transfected with plasmid and donor oligos using TranIT-LT1 (Mirus). Green fluorescence-activated cell sorting and replated in Matrigel. Single clones were manually excised for genotyping, expansion, and cryopreservation. Correctly targeted clones were identified by polymerase chain reaction (PCR), enzyme digestion, and Sanger sequencing.

### GENERATION OF DOXYCYCLINE INDUCIBLE RFX6 AND PDX1 STEM CELL LINE

For inducible overexpression, we obtained human RFX6 cDNA in pENTR221 (from ORF Clone Collection ID IOH27199), which was then cloned into lentiviral destination vector pInducer20 (gift from Stephen Elledge, Addgene no. 44010) using Gateway cloning methods with LR Clonase II (Invitrogen). Successful cloning was confirmed using Sanger sequencing. Lentiviral particles were generated by CCHMC Viral Core. iPSCs were transduced by addition of lentiviral supernatant to the culture medium immediately after passaging, which was then replaced with fresh medium after 24hrs of exposure. The cells were selected with G418 (100ug/mL) for 4 days, beginning on the following day. At this point, the colonies had normal morphology and normal characteristics, whereas mock transduced cells were dead. The cell line was the maintained under standard culture conditions, with intermittent exposure to G418 to maintain resistant cells. Inducible RFX6 expression was validated by qPCR 24hrs after exposure to doxycycline (.5ug/mL) at the pluripotent cell stage. In the same way, PDX1 inducible line was generated as previously described (Krishnamurthy et al. 2022). They were maintained in mTeSR with intermittent G418 selection.

### RNA SEQUENCING

The initial amplification step for all samples was done with the Ovation RNA-Seq System v2 kit (Tecan Genomics). The assay was used to amplify RNA samples to create double stranded cDNA. The concentrations were measured using the Qubit dsDNA BR assay. The cDNA size of each sample was determined by using an Agilent HS DNA Chip. Libraries were then created for all samples. Specifically, the Illumina protocol, the Nextera XT DNA Sample Preparation Kit, was used to create DNA library templates from the double stranded cDNA. The concentrations were measured using the Qubit dsDNA HS assay. The size of the libraries for each sample was measured using the Agilent HS DNA chip. The samples were placed in a pool. The concentration of the pool is optimized to acquire at least 40 million reads per sample. The investigator requested the use of Paired-End 100 bp Reads, so for sequencing a NovaSeq S1 (200 cycles) v1.5 flow cell was used. Libraries were sequenced on the Illumina NovaSeq 6000 system following the manufacturer’s protocol.

### BULK RNA-SEQ ANALYSIS

RNA-seq data was analyzed using Computational Suite for Bioinformaticians and Biologists (CSBB – v3.0.0) using the *Process-RNASeq_PairedEnd* and *Generate-TPM_Counts_Matrix* modules (https://github.com/praneet1988/Computational-Suite-For-Bioinformaticians-and-Biologists). FASTQs have adapters trimmed using a tool called bbduk, and quality is checked using fastqc. Bowtie2 + RSEM is used to align and count transcripts, and you should mention whatever reference genome you’re using. From the RSEM results we generate count/TPM matrices for the genes/isoforms. Further analysis and data visualization was done using RStudio v4.2.1 with the following package: DESeq2, ggplot2, plyr, ggrepel, EnhancedVolcano, pheatmap.

### IMMUNOFLUORESCENCE

Tissue was fixed in 4% paraformaldehyde, cryopreserved in 30% sucrose, embedded in OCT, and frozen at -80C cyosectioned. 8um cryosections were mounted on Superfrost Plus slides and permeabilized, blocked and stained according to standard protocol. Primary antibodies used are listed in the table below and all secondary antibodies were conjugated to Alexa Fluor 488, 546/555/568 or 647 (Invitrogen) and used at 1:500 dilution. Images were acquired using a Nikon A1 GaAsP LUNV inverted confocal microscope and NIS Elements software (NIKON). Quantification and analysis was done using NIKON software.

| *Primary antibody* | *Company* | *Host* | *Dilution* |
| --- | --- | --- | --- |
| Rfx6 | R&D AF7780 | Sheep | 1:250 |
| Chga | Abcam #ab15160 | Rabbit | 1:500 |
| Ecad | BD Transduction Lab #610182 | Mouse | 1:500 |
| Pdx1 | Abcam ab47383-100 | Goat | 1:250 |
| Foxa2 | Abnova/Novus #H00003170-M01 | Mouse | 1:500 |
| Sox2 | Millipore #AB5603 | Rabbit | 1:500 |
| Sox17 | R&D #AF1924 | Goat | 1:500 |
| Cdx2 | Cell Marque (Sigma) EPR2764Y | Mouse | 1:300 |
| Satb2 | Cell Marque EP281 | Rabbit | 1:100 |
| Ecad | R&D #AF648 | Goat | 1:500 |
| Muc2 | Cloud Clone PAA705Mu01 | Rabbit | 1:500 |
| Chga | DSHB | Mouse | 1:500 |
| Ghrl | Millipore | Rabbit | 1:250 |
| 5-HT (Serotonin) | Immunostar #20080 | Rabbit | 1:1000 |
| Defa-5 | Novus BiologicalsNB110-60002 | Mouse | 1:60 |
| GLP-1 | Abcam | Rabbit | 1:500 |
| Cdh17 | Sigma #HPA023616 | Rabbit | 1:500 |
| Pdx1 | Abcam ab47308 | Guinea pig | 1:250 |

### qPCR

RNA was extracted using the Nucleospin RNA extraction kit (Macharey-Nagel) and reverse transcribed into cDNA using Superscript VILO (Invitrogen) according to manufacturer’s instruction. qPCR primers were designed using NCBI PrimerBlast. Primer sequences are listed in the table below. qPCR was performed using Quantitect SYBR Green PCR kit (QIAGEN) and a QuantStudio 3 Flex RT PCR System (Applied Biosystems). Relative expression was determined using the Delta Delta Cycle method and normalizing to PPIA (cyclophilin A). Samples from at least three independent passages were used for quantification.

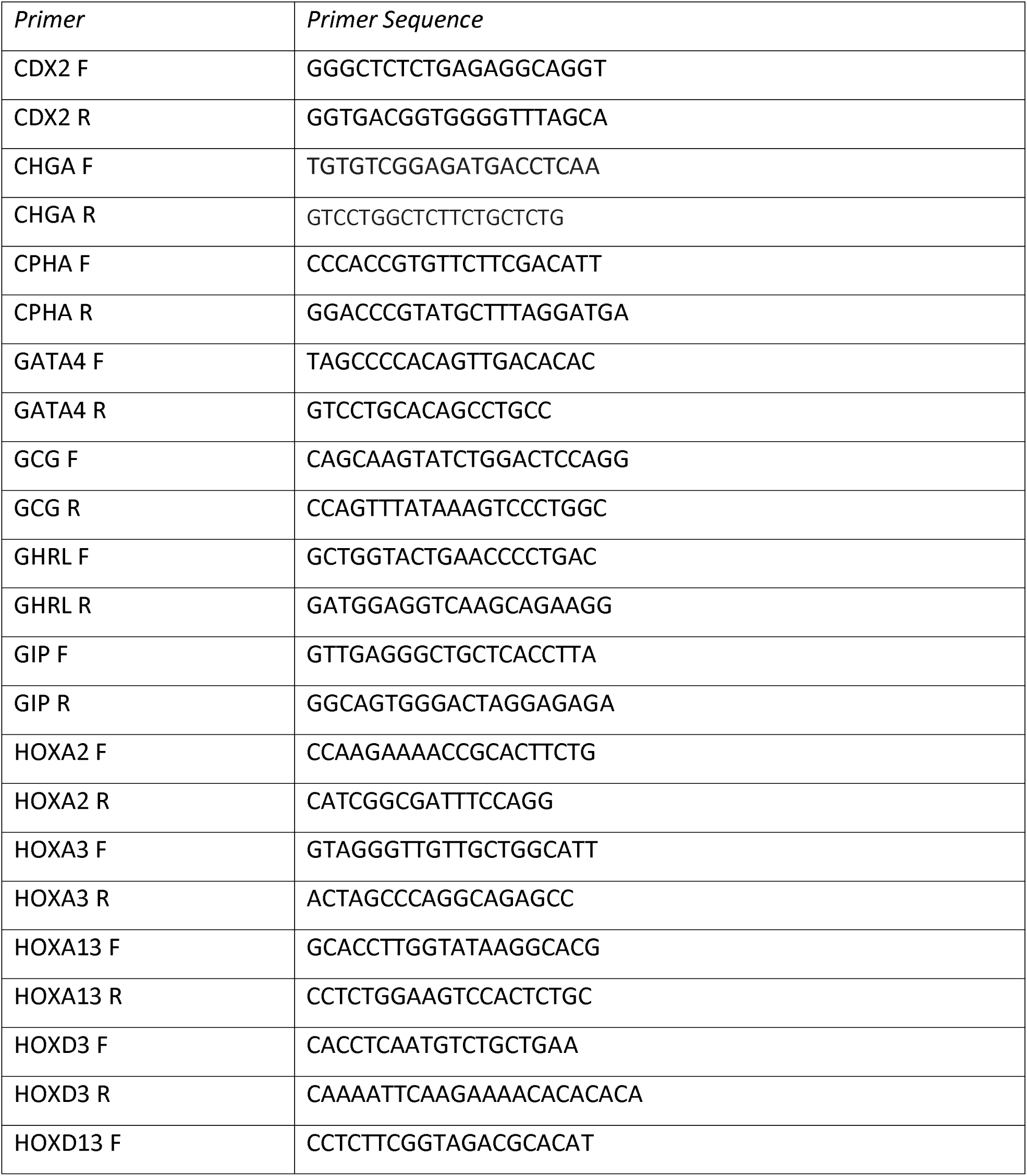
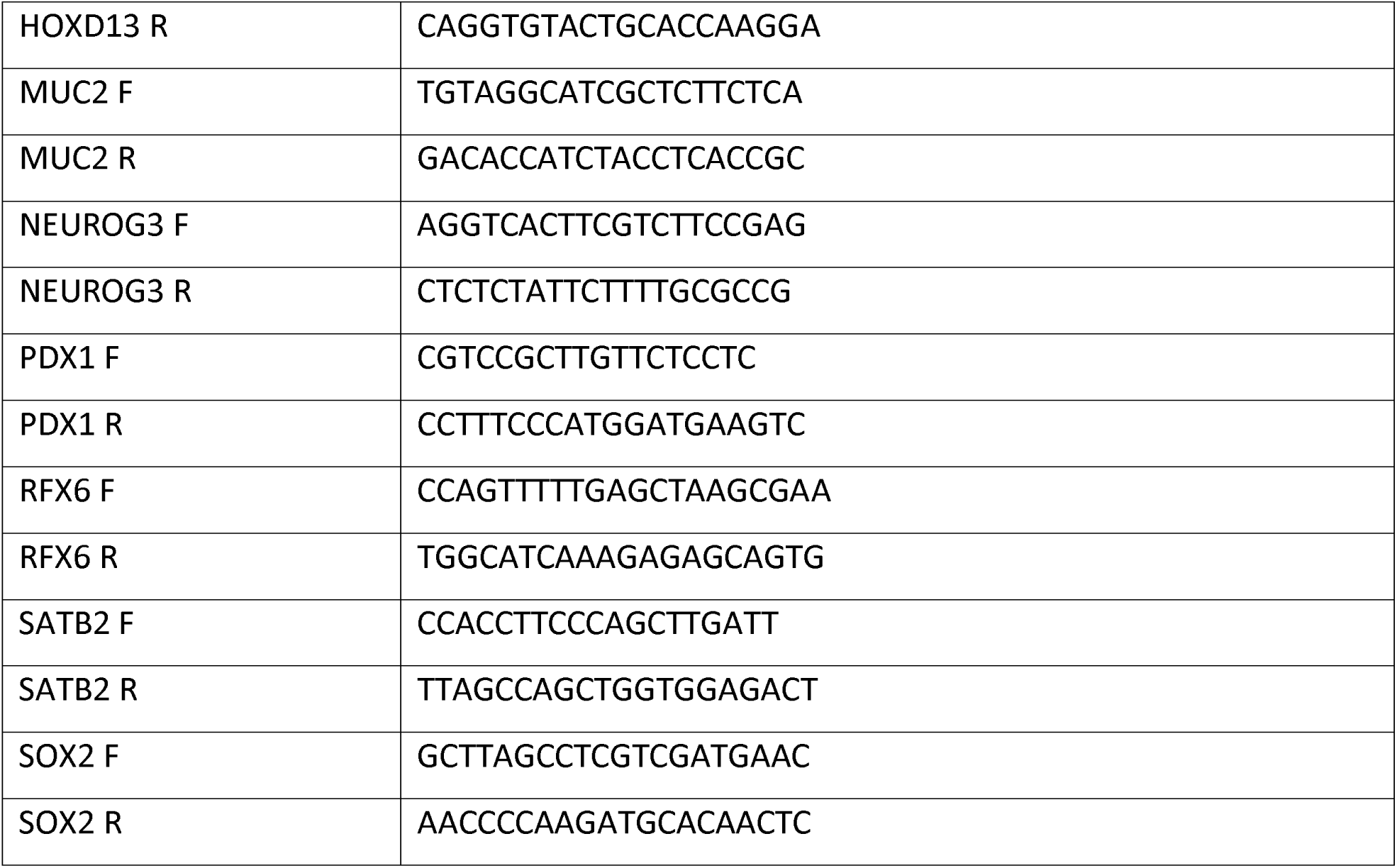

### XENOPUS METHODS

Xenopus embryos and injections: WT adult *X. laevis* frogs were purchased from the National Xenopus Resource Center(NXR, Woods Hole, MA, RRID:SCR_013731). Ovulation, in-vitro fertilization, embryo de-jellying, and microinjection were performed as described (Sive et al., 2000). Human *RFX6* cDNA clone was purchased from Horizon Discovery (OHS6084-202637733) and gateway sub-cloned from its entry vector pENTR223 into the expression vector pCSf107mT-Gateway-3⍰myc (Addgene 67617) using clonase (ThermoFisher 11791020) according to manufacturer’s instructions. SacII linearized plasmid template was used to make mRNA for injection using the Ambion mMessage mMachine SP6 RNA Synthesis Kit (ThermoFisher AM1340) according to manufacturer’s instructions.

Wild type (wt) human *RFX6* coding region (wt: NCBI Reference Sequence: NM_173560.4) and mutated versions (a 1bp mutation nucleotide C2623T or a 13bp deletion nucleotides 1040-1052 of the coding sequence) were also commercially synthesized with 5’EcoRV and 3’SpeI ends and cloned into the pCDNA3.1 vector by Genscript (Genscript USA, Piscataway, NJ). To construct the inducible GR-*RFX6* wild type or mutants used in cycloheximide direct target assays, the respective *RFX6* wt or mutant versions were released from pCDNA3.1 via EcoRV/SpeI digests, gel purified, and cloned 5’EcoRV 3’SpeI into the pT7TS-GR-HA vector. XbaI linearized plasmid templates were used to make mRNA for injection using the Ambion mMessage mMachine T7 mRNA Synthesis Kit (ThermoFisher AM1344) according to manufacturer’s instructions. 100pg total mRNA for wt or mutant GR-*RFX6* RNAs were injected at the 8-cell stage into ventral-vegetal blastomeres to target the hindgut (50pg per cell). At stage 20, hindgut explants (containing both endoderm and mesoderm) were micro-dissected in 1× MBS +50 μg/ml gentamycin sulfate (gent; MP Biochemicals 1676045) and cultured in 0.5× MBS +0.2% fatty acid free BSA (Fisher BP9704) +50 μg/ml gent with the following concentrations of factors: 1 μM dexamethasone (DEX; Sigma D4902); 1 μM cycloheximide (CHX; Sigma C4859). In CHX experiments, explants were treated for 2 hr in CHX prior to DEX+CHX treatment for 6 hr.

The previously validated translation-blocking morpholino oligo (MO) against Xenopus Rfx6 (Pearl et al 2011) was purchased from GeneTools (Philmath, OR) and injected at the 8-cell stage (1.5ng per cell) into each dorsal vegetal blastomere to target the foregut. MO sequences were as follows: Rfx6-MO1: 5⍰ AAT TGG CAT TTC ACC GGG TTC AGG C 3⍰; a negative control Rfx6 mismatch (MM MO) with five bases altered: MM MO: 5⍰ AAT **<u>a</u>**GG **<u>g</u>**AT TT**<u>g</u>** ACC **<u>c</u>**GG TTC A**<u>c</u>**G C 3⍰ (mismatch bases indicated in lowercase, bold, underlined).

Xenopus RT-qPCR: Xenopus explants were dissected from embryos of 2–3 separate fertilization/injection experiments, homogenized and frozen on dry ice in 350 μl of Buffer RA1 containing BME (Macherey-Nagel Nucleospin RNA isolation kit 740955), and stored at –80°C. RNA was extracted according to manufacturer’s instructions and 500ng RNA was used in cDNA synthesis reactions using Superscript Vilo Mastermix (ThermoFisher 11755050), also according to the manufacturer’s instructions. qPCR reactions were carried out using PowerUp Mastermix (ThermoFisher A25742) on an ABI QuantStudio3 machine. *Xenopus* RT-qPCR primer sequences were as follows: *pdx1.L* forward: 5’GTT CCC TCA GCT GCT TAT CG; *pdx1.L* reverse: 5’TAC CAA GGG GTT GCT GTA GG; *ptf1a.L* forward: 5’ATG GAA ACG GTC CTG GAG CA; *ptf1a.L* reverse: 5’GAG GAT GAG AAG GAG AAG TTG. Relative expression, normalized to ubiquitously expressed *odc*, and then set relative to uninjected, untreated hindgut explants, was determined using the 2^−ΔΔCt^ method. Graphs display the average 2^−ΔΔCt^ value ± standard deviation. Statistical significance (p<0.05) was determined using parametric two-tailed paired t-test, relative to uninjected, untreated hindgut explants. Each black dot in the RT-qPCR graphs represents an independent biological replicate containing four explants.

*Xenopus* in-situ hybridization:In-situ hybridization of *Xenopus* embryos was performed as described (Sive et al., 2000) with minor modifications. Briefly, embryos were fixed overnight at 4°C in MEMFA (0.1 M MOPS, 2 mM EGTA, 1 mM MgSO4, and 3.7% formaldehyde), washed 2×10 min in MEMFA without formaldehyde, dehydrated directly into 100% ethanol, washed 4– 6 times in 100% ethanol, and stored at −20°C in 100% ethanol for at least 24 hr. Proteinase K (ThermoFisher AM2548) on day 1 was used at 2 µg/ml for 10 min on stage 20 embryos and 5 µg/ml for 10 min on NF33 embryos; hybridization buffer included 0.1% SDS; the RNAse step was omitted; and anti-DIG-alkaline phosphatase antibody (Sigma 11093274910) used at 1:5000 in MAB buffer (100 mM Maleic acid, 150 mM NaCl, and pH 7.5) + 20% heat-inactivated lamb serum (Gibco 16070096) + 2% blocking reagent (Sigma 11096176001). Full length*rfx6.L* and *pdx1.L* anti-sense DIG-labeled in-situ probes were generated using linearized plasmid cDNA templates with 10× DIG RNA labeling mix (Sigma 11277073910) according to the manufacturer’s instructions.

Xenopus luciferase assays: A 1,236 bp sequence of the human *PDX1* upstream “area I II III” enhancer (>hg19_dna range=chr13:28491064-28492299) was screened for RFX6 responsiveness due to the observation that the orthologous mouse *Pdx1* enhancer region has previously been shown to drive reporter expression in foregut and duodenum (Gannon et al 2001; Fujitana et al 2006). Wild type or a mutant version of the human *PDX1* enhancer with 6 CisBP predicted RFX binding motifs mutated (Weirach et al 2014) were commercially synthesized by Genscript (Genscript USA, Piscataway, NJ) and cloned into the pGL4.23 firefly luc2/miniP vector (Promega E8411). 16/32-cell embryos were co-injected with 5 pg of pRL-TK:renilla luciferase plasmid (Promega E2241) + 80 pg of the pGL4.23 luc2/miniP Human*PDX1* enhancer:firefly luciferase plasmid (wt or 6RFX motif mutant) into either a D1 blastomere to target foregut and a D4 blastomere to target hindgut. The following amounts of MOs or mRNAs were also injected in enhancer:luciferase assays: 3ng total of Rfx6-MO1 or 5bp mismatch-MO; 100pg wt or mutant human *RFX6* mRNAs.

Each biological replicate contained a pool of five embryos, obtained from 2 to 3 separate fertilization/injection experiments, which were frozen on dry ice in a minimal volume of 0.1× MBS and stored at –80°C. To assay luciferase activity, samples were lysed in 100 µl of 100 mM TRIS-Cl pH 7.5, centrifuged for 10 min at ∼13,000×*g* and then 50 µl of the clear supernatant lysate was used separately in firefly (Biotium #30085-1) and renilla (Biotium 300821) luciferase assays according to the manufacturer’s instructions. Relative luciferase activity was determined by normalizing firefly to renilla levels for each sample. Graph show the average relative luciferase activity ± standard deviation with black dots showing values of biological replicates. Statistical significance was determined by parametric two-tailed paired t-test, *p<0.05.

### DATA AVAILABILITY

Raw data and processed files have been submitted to the GEO repository. Accession number: GSE245474

### STATISTICS

Data are presented as the mean +/- standard error of the mean. Significance was determined using appropriate tests in Graph Pad Prism, with P>.05 not significant; *P<.05. **P<.01, ***P<.001.

## AUTHOR CONTRIBUTIONS

Conceptualization: JGS, JMW. Methodology: JGS, SAR, HAM, EFP, DOK, JER, MK, AZ, JMW. Formal analysis: JGS, SAR, EFP. Investigation: JGS, SAR. Resources (biopsytissue and patient recruitment): MK, NAH, SAG, LLF. Writing of original draft: JGS, JMW. Editing: JGS, SAR, HAM, DOK, JMW. Visualization: JGS. Supervision: JMW. Project administration: JMW. Funding acquisition: JMW.

## ACKNOWLEDGEMENTS

We thank the support provided by the confocal imaging core, gene expression core, pathology core and Pluripotent stem cell facility at CCHMC. We thank all the members of the Wells and Zorn labs for reagents and feed back. All figures were created with Adobe illustrator, BioRender.com and Graphpad-Prism.

## Figure Legends

**Fig S1-Manipulating exposure of growth factors to pattern proximal and distal HIOs**A) Schematic of HIO differentiation with modified protocol to obtain either Duodenal or Ileal HIOs. B) Changes in Duod and Ile HIOs in patterning markers (CDX2, PDX1, SATB2) (n=4-8) (scale bar=100um). C) Enteroendocrine cells presence in both Duod and Ile HIOs (CHGA) as well as different Populations of EECs found in Duo vs Ile HIOs forming EEC-secreted hormone gradient seen in the human GI tract (scale bar =100um). D) Heatmap comparing significantly differentially expressed genes in Duo and Ile HIOs (n=3). E) Volcano Plot showing differentially expressed genes between Duod and Ile HIOs. Significant Differentially expressed genes from Bulk RNA sequencing was defined by adjusted p-value of <.05.

**Fig s2-Rfx6 expression throughout the intestinal differentiation** A) Schematic of generation of iPSC from patient sample and HIOs from patient iPSC. B) Brightfield images of HIOs at D7, D35 and after transplant into NSG mouse kidney capsule (10wks). C) Expression of RFX6 throughout intestinal differentiation at definitive endoderm (DE), D5 and D7 hindgut patterning stages and *in vitro* end stage (D35). D) Staining of RFX6 expression at DE, D5 and D7 monolayers of HIO differentiation (n=5). Significance determined by unpaired t-test with *p<0.05, **p<0.01, ***p<0.001.

**Fig S3-Rfx6 mutation affects differentiation of enteroendocrine cells** A) Patient biopsy staining for enteroendocrine subpopulations and phenotypes in Rfx6 Mut Patient, loss of GLP1 lineage and increase in 5HT with the quantification (n=3-8). B) Rfx6 Mut HIOs replicate endocrine phenotypes seen in biopsy. C) Additional enteroendocrine subtype lineages lost in Rfx6 Mut HIOs (Ghrl, Motilin and GIP). D) Schematic of hypothesized Rfx6 place in EEC differentiation gene regulatory network.

**FigS4 – Transcriptomic changes in RFX6 Mutant** in vivo A) PCA of WT vs RFX6 Mut Duod HIOs after transplant B) Volcano plot and heatmap of transplanted HIOs WT vs Rfx6 Mut Significantly Differentially Expressed Genes (n=3). C) Distal Hox genes expression in WT HIOs vs RFx6 Mut showing an increase in RFX6 mut transplanted HIOs. D) PCA of WT vs RFX6 Mut and CRISPR Corrected transplanted HIOs. E) Heatmap of differentially expressed genes (n=3). F) Proximal small intestine patterning markers, and duodenal functional genes are upregulated after correction of paternal allele while distal patterning markers decrease in CRISPR Corrected HIOs. Significance determined by unpaired t-test with *p<0.05, **p<0.01, ***p<0.001. Significant Differentially expressed genes from Bulk RNA sequencing was defined by adjusted p-value of <.05.

**Fig S5-RFX6 Mut D7 HIOs shows mis-patterning of developing hindgut** A) Volcano plot of significantly differentially expressed genes between WT and RFX6 Mut D7 HIOs. B) CDX2 is downregulated in RFX6 Mut D7 HIOs and known CDX2 targets are also downregulated (n=4). C) Overlap of significantly downregulated genes in RFX6 Mut D7 HIO and Endoderm markers from published dataset (Han et al) and GO term analysis for top biological process of overlapped genes. D) Overlap of downregulated genes in RFX6 Mut D7 HIO and Mesoderm markers from published dataset (Han et al) and GO term analysis for top biological process of overlapped genes. E) Volcano plot of significantly differentially expressed genes in WT vs Mut RFX6-CRISPR Corrected D7 HIOs F). Volcano plot of significantly differentially expressed genes in Mut vs Mut RFX6-CRISPR Corrected D7 HIOs. Significance determined by unpaired t-test with *p<0.05, **p<0.01, ***p<0.001. Significant Differentially expressed genes from Bulk RNA sequencing was defined by adjusted p-value of <.05.

**Fig S6-RFX6 regulates pdx1 expression in Xenopus developing foregut.** A) In situ hybridization of *rfx6* (expression domain abbreviations: st, stomach; duo, duodenum; vp, ventral pancreas; dp, dorsal pancreas; gb, gall bladder) and *satb2* (abbreviation: hg, hindgut) in stage NF33 (2.5 d.p.f) Xenopus embryos. Numbers in the lower left corner of panel indicate number of embryos assayed with the shown gene expression pattern. B) Knockdown of Rfx6 in Xenopus using translation-blocking morpholino at NF33 and microdissection of the gut tube at NF43. In situ hybridization of *pdx1* (expression domain abbreviations: st, stomach; duo, duodenum) C) Positive control for CHX effectiveness, expression of the pancreas TF ptf1a. D) pdx1 enhancer luciferase activity in the HG by misexpression of WT or mutant RFX6 mRNA at different doses.

**Fig S7-Inducible constructs were built and used during the HIO differentiation.**A) Timeline for induction of Pdx1 and Rfx6 via Doxycycline throughout the intestinal differentiation. B) Schematic of doxycycline-inducible construct used in HIO differentiation. C) Overlap of genes downregulated genes in RFX6-Mut D7 HIOs and Upregulated in RFX6-Mut PDX1 Inducible genes showing a rescue of most of the genes. D) Overlap of genes downregulated genes in RFX6-Mut D7 HIOs and Upregulated in RFX6-Mut RFX6 Inducible genes. E) Differentially expressed between RFX6-Mut RFX6 Inducible and RFX6-Mut PDX1 Inducible D7 HIOs showing RFX6 induction upregulates more genes than PDX1 in the RFX6 Mut HIOs. F) GO term analysis for Biological Processes and Molecular Function of PDX1 induction vs RFX6 induction in RFX6 Mut HIOs. Significant Differentially expressed genes from Bulk RNA sequencing was defined by adjusted p-value of <.05.

**TableS1-Alignments of the human/mouse/frog conserved pdx1 enhancer with the 6 RFX motifs highlighted.**

## Notes

### Competing Interest Statement

The authors have declared no competing interest.

